# Targeted Long-Read RNA Sequencing Demonstrates Transcriptional Diversity Driven by Splice-Site Variation in *MYBPC3*

**DOI:** 10.1101/522698

**Authors:** Alexandra Dainis, Elizabeth Tseng, Tyson A. Clark, Ting Hon, Matthew Wheeler, Euan Ashley

## Abstract

**Background:** Clinical sequencing has traditionally focused on genomic DNA through the use of targeted panels and exome sequencing, rather than investigating the potential transcriptomic consequences of disease-associated variants. RNA sequencing has recently been shown to be an effective additional tool for identifying disease-causing variants. We here use targeted long-read genome and transcriptome sequencing to efficiently and economically identify molecular consequences of a rare, disease-associated variant in hypertrophic cardiomyopathy (HCM).

**Methods and Results:** Our study, which employed both Pacific Biosciences SMRT sequencing and Oxford Nanopore Technologies MinION sequencing, as well as two RNA targeting strategies, identified alternatively-spliced isoforms that resulted from a splice-site variant containing allele in HCM. These included a predicted in-frame exon-skipping event, as well as an abundance of additional isoforms with unexpected intron-inclusion, exon-extension, and pseudo-exon events. The use of long-read RNA sequencing allowed us to not only investigate full length alternatively-spliced transcripts but also to phase them back to the variant-containing allele.

**Conclusions:** We suggest that targeted, long-read RNA sequencing in conjunction with genome sequencing may provide additional molecular evidence of disease for rare or de novo variants in cardiovascular disease, as well as providing new information about the consequence of these variants on downstream RNA and protein expression.

## INTRODUCTION

As clinical genome sequencing becomes more widespread, we continue to gain more information about the genetic basis of disease. Large cohort studies provide insight into the myriad genetic variants that can have combinatorial effects on common diseases while individual sequencing can reveal rare or de novo variants in mendelian disease. Genetic testing can provide immediate value to patients and their families by clarifying diagnoses, directing therapy, and facilitating cascade screening.

To date, clinical sequencing has focused mostly on examination of genomic DNA using targeted panels and exome sequencing. However, recent sequencing of a large hypertrophic cardiomyopathy (HCM) cohort revealed that positive identification of a disease-associated variant was returned in only 32% of patients, with an additional 15% resulting in an inconclusive result^1^. Recent studies have suggested that turning to whole genome sequencing rather than targeted panels may improve the yield of finding disease-associated variants^2^. One such example used whole genome sequencing to identify four novel, deep-intronic mutations in *MYBPC3* with the potential to modify splicing in a cohort of patients. Yet when RNA from each patient was examined, only three of the four mutations actually resulted in improper splicing, highlighting the need for RNA based assays to confirm potential effects of novel mutations before assigning them disease-association or causality^2^.

In patients where genome sequencing fails to reveal causative variants, or where patients are left with variants of unknown significance (VUS), sequencing of the transcriptome may also provide additional diagnostic clarity. Transcriptome analysis has revealed or confirmed causative, non-coding mutations not previously found in clinical sequencing in a number of diseases, including muscular dystrophy and a novel neuromuscular disease^3,4^. A recent cohort study examining 50 patients with genetically undiagnosed muscle disorders found that RNA-Sequencing, when used as a complementary tool to exome and whole genome sequencing, had an overall diagnosis rate of 35%^5^. There is mounting evidence that investigating the transcriptome in addition to the genome may yield additional insight when rare or de novo mutations are found, including non-coding mutations. Here, we demonstrate the clinical application of a long-read, targeted transcriptome sequencing strategy.

Traditional next generation sequencing has consisted of short-read sequencing, which sequences short fragments of DNA, typically between 75-300bp in length. While short read sequencing is an efficient and economic method of whole genome sequencing, long-range genomic information, including phasing of variants and large structural variation, can be lost. As an alternative, new long-read sequencing technologies, including Pacific Biosciences SMRT Sequencing and Oxford Nanopore Technologies MinION Sequencing, which can sequence DNA in fragments of tens to hundreds of kilobases in length, retains this information. For genomic analysis, this can provide valuable information about the phase of disease causing variants. In RNA analysis, long reads can provide complete pictures of the splicing occurring in full-length transcripts, rather than focusing only on individual splice junctions, and can provide phased information linking an improperly spliced transcript back to a disease-associated allele.

Though rich in long-range information, whole genome or whole transcriptome long-read sequencing to a clinically relevant depth can become cost inefficient. While prices continue to fall, targeted sequencing strategies, which focus sequencing capacity only on genes of interest, can provide a current solution to sequencing disease-relevant genes across multiple patients. Targeting strategies may include PCR-primer based methods^6^, probe based pulldown^7–9^, or even more recent Cas9-targeted sequence capture^10^, but all strive to increase the efficiency of long-read sequencing methodologies when interested in a defined set of targets. For our investigation, we were interested in a small number of transcriptomic and genomic targets associated with HCM.

HCM, characterized by an overgrowth of the left ventricular wall, affects 1:500 in the population and can lead to atrial fibrillation, heart failure, and sudden death^11^. Causative genetic mutations can be identified in 30-60% of sequenced patients, and 75% of those appear on one of two genes: *MYH7* (myosin heavy chain 7) or *MYBPC3* (cardiac myosin binding protein c), key components of the cardiac sarcomere^12,13^.

The majority of these disease associated variants in *MYBPC3* are nonsense or frameshift mutations, unlike other HCM-associated genes, including *MYH7*, where missense mutations account for the majority of disease-causing variants^14^. Haploinsufficiency is therefore thought to be a major contributor to the pathogenicity of disease-associated *MYBPC3* alleles and appropriate expression levels have been shown to be critical during development^15^. Mutations in *MYBPC3* that affect splice sites have additionally been shown to be linked to HCM, though many reported mutations have been predicted to result in a frameshift, likely resulting in a premature stop codon, nonsense-mediated decay, and haploinsufficiency as described above^16^. One of the first *MYBPC3* mutations linked to HCM was a splice acceptor mutation just upstream of exon 21 that induced skipping of this exon, though this mutation was predicted to cause disease via haploinsufficiency, as it also resulted in a premature stop codon^17^.

We here investigate a putative splice-site-altering mutation in *MYBPC3* found in a 21-year-old female patient with hypertrophic cardiomyopathy. A sequencing panel identified a single-base pair change in *MYBPC3* (c.1898-1G>A) that was predicted to disrupt the exon 20 splice-acceptor site. We used targeted long-read sequencing to sequence full-length *MYBPC3* transcripts and revealed a diverse set of alternatively-spliced isoforms that resulted as a consequence of this mutation. We additionally used targeted long-read genomic sequencing to assign the alternatively spliced isoforms to the mutation-containing allele. This manuscript demonstrates the ability to use commercially available target-capture probes for not just Pacific Biosciences SMRT sequencing but also Oxford Nanopore MinION sequencing of both genomic and transcriptomic targets. It additionally reveals transcript level consequences of a rare, uncharacterized variant in *MYBPC3*. Together, these points demonstrate the potential of targeted, long-read RNA sequencing to be an effective diagnostic tool when faced with rare or de novo mutations.

## METHODS

### Patient Phenotyping and Clinical Genetic Testing

We here investigate a predicted splice-site-altering mutation in *MYBPC3* which presented in a 21-year-old female patient with hypertrophic cardiomyopathy. The patient first presented with a heart murmur and HCM at 13 years old, and had an implantable cardioverter-defibrillator (ICD) placed at 16 due to a septal wall thickness of 3.5 cm. Upon evaluation at age 21, the patient was experiencing palpitations and symptoms consistent with New York Heart Association Functional Class III status. An echocardiogram showed systolic anterior motion of the mitral valve and severe concentric left ventricular hypertrophy with an intracavitary gradient with severe bi-atrial enlargement and an ejection fraction of 77%. The patient underwent a myectomy to remove obstructive myocardial tissue, which was then later used for RNA extraction.

An HCM panel (GeneDx, Gaithersburg, MD) identified a single-base pair change in *MYBPC3* (c.1898-1G>A) that was predicted to disrupt the exon 19-20 splice junction. This mutation disrupts the splice acceptor site just before exon 20 and is predicted to result in a broken WT splice acceptor site by the Human Splicing Finder tool^18^. This is likely to result in skipping of exon 20, with exon 19 spliced directly to exon 21. Exon 20 composes part of a short linker region between the C4 and C5 immunoglobulin-like domains of *MYBPC3^19^*. This mutation has been previously noted in one patient from a cohort of 200 unrelated Chinese adult HCM patients^20^ and is not present in the Genome Aggregation Database (gnomAD) as of December 2018^21^.

### Preliminary PacBio Iso-Seq of *MYBPC3*

We created a multiplexed, barcoded PacBio Iso-Seq library to sequence full-length *MYBPC3* cDNA in our study patient sample as well as 9 additional samples [Figure 1a]. These included six healthy control hearts and three hearts with known HCM-associated point mutations in *MYH7*. Two of the point-mutation samples and the *MYBPC3* splice-variant sample were collected from septal myectomies while the remaining samples, including all controls, were from explanted hearts. Hearts were collected under Stanford Institutional Review Board GAP approval number 4237.

**Figure 1:**
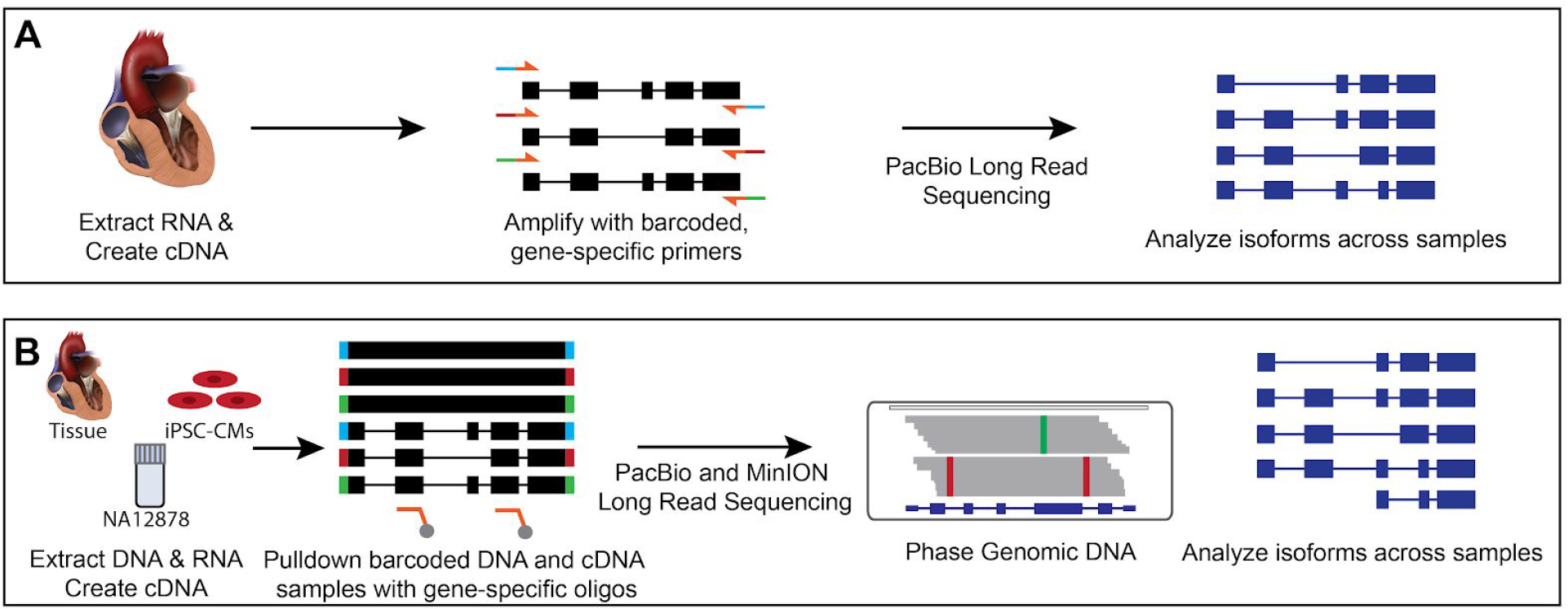
Strategies for investigating alternative splicing and phasing in cardiovascular disease genes. In our first method of phasing cDNA from left ventricular (LV) heart samples (Panel A) we created cDNA from RNA extracted from frozen LV samples and amplified a transcript of interest (*MYBPC3*) with barcoded primers specific to the first and last exons. We pooled barcoded samples and sequenced using Pacific Biosciences SMRT sequencing. This allowed us to phase and compare isoforms across samples. In our second strategy (Panel B), we extracted DNA and RNA from the same samples, which included NA12878, an iPSC-CM cell line, and a left ventricular heart sample. We then created cDNA from the RNA, and barcoded it as well as fragmented genomic DNA. We pulled both down using IDT xGen oligos specific to the exons of ten cardiovascular genes of interest. We pooled our samples and sequenced using either PacBio’s SMRT sequencing or Oxford Nanopore Technologies’ MinION sequencing. This allowed us to phase the genomic DNA as well as the cDNA, and analyze isoforms across samples.

RNA was extracted from hearts using Qiagen’s miRNeasy kit. First strand cDNA synthesis was performed using Superscript III RT. Amplification was then performed using Primestar GXL and barcoded primers designed to target the first and last exon of *MYBPC3* [Supplemental Table 1]. Amplified isoforms were pooled such that each pool contained *MYBPC3* (and an additionally tested gene, *MYH7*) transcripts from five samples. These two resulting pools were turned into PacBio libraries using the standard damage repair and adapter ligation protocol. Samples were sequenced on an RSII using binding kit P6-C4 Chemistry. Each pool was sequenced on 4 SMRT cells using 4 hour movies. Sequencing was performed using a Pacific Biosciences RSII at the Stanford Genome Sequencing Service Center.

After sequencing, isoforms were mapped to hg19 (GMAP version 2014-10-22)^22^ and filtered for ≥ 99% coverage and ≥ 95% identity. Isoforms with read count support < 5 were discarded. Isoform sequences were required to begin and end with designed primer sequences. *MYH7* and *MYBPC3* isoforms were chained across all samples to find common isoforms and look for patterns in isoform expression.

### Hybridization and Pulldown of Genomic Libraries for Targeted Sequencing

In addition to targeted genomic sequencing of our patient sample on both the Pacific Biosciences and Oxford Nanopore platforms, we sequenced two other samples to act as controls: NA12878, a well-studied benchmarking sample for genome sequencing technologies, and iPSC-cardiomyocytes containing an HCM-associated point mutation, R403Q, in *MYH7* (R403Q iPSC-CMs, Stanford Institutional Review Board GAP approval number 4237). The following cell lines/DNA samples were obtained from the NIGMS Human Genetic Cell Repository at the Coriell Institute for Medical Research: NA12878, GM12878.

DNA was extracted from 11mg flash-frozen myectomy tissue from our case study patient and the iPSC-CMs using Qiagen’s DNeasy Blood and Tissue spin kit. DNA from NA12878 was extracted by Coriell. DNA was sheared using a Covaris g-TUBE in an Eppendorf 5415C centrifuge at 6000 RPM. AMPure XP bead purified DNA was then size selected for 6-13kb fragments on a BluePippin (Sage Science). Barcoded adapters (From PacBio Protocol “Target Sequence Capture Using SeqCap EZ Libraries with PacBio Barcoded Adapters.” ^23^) were ligated using KAPA’s Hyper Prep Kit and DNA was amplified using Hot-Start LA Taq DNA Polymerase from Clontech for a total of 9 PCR cycles, using Pacbio’s universal oligos as primers (from “Target Sequence Capture Using SeqCap EZ Libraries with PacBio Barcoded Adapters.” ^23^).

Amplified DNA from 3 samples was pooled before pulldown with IDT xGen probes designed against a panel of 10 cardiac genes of clinical interest: *LMNA*, *MYBPC3*, *MYH6*, *MYH7*, *MYL2*, *PKP2*, *RYR2*, *TNNT2*, *TPM1*, & *TTN*. Pulldown was performed for 4 hours at 65C, and probes were captured after pulldown using M-270 streptavidin beads and washed using IDT’s xGen Lockdown Reagents. Captured samples were then PCR amplified again using LA Taq for a total of 18 cycles (20 cycles for R403Q sample) using PacBio’s Universal Oligos. Samples were size-selected again for 6-13.5kb fragments, and then amplified for 4 more cycles.

Samples were then split into two pools: one for PacBio SMRT sequencing and one for Oxford Nanopore MinION sequencing. PacBio SMRTbell libraries were prepared using PacBio’s standard library preparation kit and protocol. MinION libraries were prepared for 1D^2 sequencing using kit SQK-LSK308.

### Hybridization and Pulldown of Transcriptomic Libraries for Secondary Sequencing

RNA was extracted from GM12878 LCLs (Coriell Institute) and R403Q iPSC-CMs using a trizol-chloroform extraction. RNA from study patient heart sample (39mg) was also extracted using a trizol-chloroform extraction, but sample homogenization was performed in trizol using a glass dounce homogenizer on ice. RNA that had been previously extracted from this sample to do the preliminary Iso-Seq above was also included as a fourth sample. First strand cDNA was created using the Clontech SMARTer cDNA synthesis kit and amplification of cDNA was performed for 14-18 cycles based on per-sample optimization. BluePippin size-selection was performed for 3000-9000bp transcripts. This would include major isoforms of *LMNA*, *MYBPC3*, *MYH7*, and *PKP2* from our cardiac gene panel. Barcoded adapters (as described above) were ligated using KAPA’s Hyper Prep Kit and DNA was amplified using Hot-Start LA Taq DNA Polymerase from Clontech for a total of 9 PCR cycles, using PacBio’s universal oligos as primers.

Amplified cDNA from 4 samples (GM12878, R403Q iPSC-CMs, and two separate RNA extractions of *MYBPC3* study patient) was pooled before pulldown with IDT xGen probes designed against a panel of 10 cardiac genes of clinical interest: *LMNA*, *MYBPC3*, *MYH6*, *MYH7*, *MYL2*, *PKP2*, *RYR2*, *TNNT2*, *TPM1*, *TTN*. Pulldown was performed for 4 hours at 65C, and probes were captured after pulldown using M-270 streptavidin beads and washed using SeqCap EZ Hybridization and Wash kit. Captured samples were then PCR amplified again using LA Taq for a total of 15 cycles using PacBio’s Universal Oligos. Samples were then split into two pools: one for PacBio SMRT sequencing and one for Oxford Nanopore MinION sequencing. SMRTbell libraries were prepared using PacBio’s standard library preparation kit and protocol. Final PacBio libraries were size-selected once again on the BluePippin to remove all fragments below 3000bp. MinION libraries were prepared for 1D^2 sequencing using kit SQK-LSK308.

### Sequencing and Data Analysis

Pacbio SMRTbell libraries were pooled and sequenced on two PacBio Sequel SMRT cells, one for the genomic libraries and one for the Iso-Seq transcriptomic libraries (SMRT Cell 1M v2 LR cells, 20 hour movies). Sequencing was performed by the DNA Sequencing & Genotyping Center at the University of Delaware.

Circular Consensus Sequences (CCS) from both the genomic and IsoSeq runs were called using SMRTLink v5.1.0, with parameters set to require at least 2 reads of insert for the Genomic library and at least 1 read for the IsoSeq library. Genomic data was converted to FASTQ formats using SMRTLink and demultiplexed using Porechop^24^. Reads were aligned to hg38 using minimap2 (version 2.10)^25^.

MinION libraries were prepared using Oxford Nanopore’s 1D^2 sequencing kit and sequenced on two R9.5 flowcells. One flowcell was used to sequence exclusively the patient genomic pulldown sample while the other flowcell was used to sequence a pooled library of genomic and transcriptomic pulldowns from all three samples. 1.28ug of starting material was used for 1D^2 library prep (ONT kit SQK-LSK308) for the patient genomic sample. 1ug of starting material was used for the same kit for the pooled samples. cDNA comprised only 10% of the pooled sample to minimize potential mispairing of closely related cDNA amplicons during 1D^2 basecalling.

MinION data analysis was performed by first using Albacore v.2.2.7 to basecall 1D and 1D^2 reads. Pooled libraries were demultiplexed using Porechop^24^. Both genomic and transcriptomic reads were aligned to hg38 using minimap2 (version 2.10)^25^.

Alternative splicing patterns in *MYBPC3* were manually identified in the Integrated Genomics Viewer (IGV)^26^, and a reference list of isoforms was created. For the purpose of this analysis, we focused on alternative splicing in the Exon 19-21 region when creating these isoforms for alignment. A small amount of alternative splicing that happens both up and downstream in both control samples and our patient samples was ignored to allow focus on splice events directly related to the *MYBPC3* splice-site mutation. Demultiplexed sequencing data was remapped to the list of isoforms using minimap2 (version 2.10) to analyze prevalence of each alternative splicing pattern in the sample.

## RESULTS

### Iso-Seq Using Gene-Specific Primers

64 high confidence isoforms were output for *MYBPC3* across 10 sequenced samples. Isoforms were “chained” across all samples to find common isoforms and look for patterns in isoform expression. Chaining of *MYBPC3* isoforms resulted in one dominant isoform per gene across all ten samples [Supplementary Figure 1].

Our study patient sample showed an increase in the number of highly expressed alternate isoforms in *MYBPC3*. In addition, it showed one highly expressed, unique alternate isoform (AS 1.1, Figure 2a). This isoform was missing exon 20, with exon 19 spliced directly to exon 21 [Figure 2b], as predicted from the (c.1898-1G>A) mutation which disrupts the splice acceptor site directly before exon 20. About 46% of the aligned reads showed improper splicing around exon 20.

**Figure 2:**
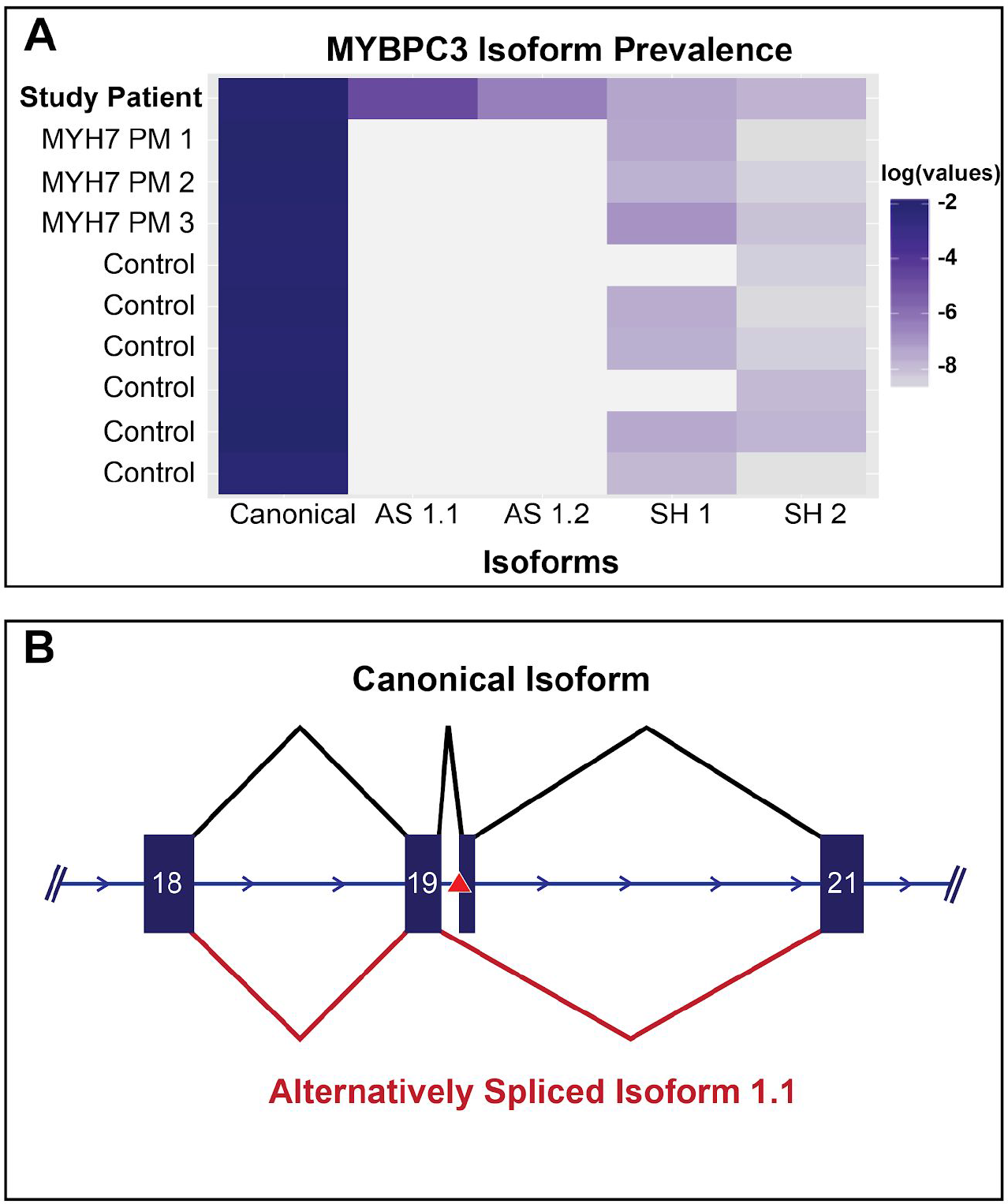
Study Patient Sample Contains Private Alternatively Spliced Isoforms. Targeted sequencing of *MYBPC3* cDNA in ten left ventricular heart samples: six healthy heart controls, our *MYBPC3* splice-variant study patient, and three HCM samples containing *MYH7* point mutations (*MYH7* PM 1-3). Sequencing revealed that our study patient, whose genome contained a splice-site variant in *MYBPC3*, contained a highly expressed alternatively spliced isoform. Panel A shows IsoSeq normalized expression (number of full length reads per isoform over the total number of full length reads) of the top 5 highly expressed *MYBPC3* isoforms in our study patient identified by PacBio’s IsoSeq pipeline across all ten samples. The canonical isoform is the most highly expressed isoform across all ten samples, but the next two highly expressed isoforms (Alternatively Spliced 1.1 and 1.2, AS1.1 and AS1.2 respectively) are seen only in our study patient sample. SH 1 and SH 2 depict shared alternatively spliced isoforms across both control and disease samples. AS1.1, the next most highly expressed isoform, is depicted in Panel B, and results in the skipping of exon 20. The location of the splice-site variant (located one base prior to the start of exon 20) is denoted with a red triangle.

As the splice-site variant seen in our patient resides in an intronic region, we were unable to definitively phase the alternatively spliced isoform to either the wild-type or variant-containing genomic allele. We were, however, able to detect that reads supporting the alternatively spliced isoform appeared to be originating mainly from the same allele. In order to phase the alternative isoform with the splicing variant, we created long-read sequencing libraries of both DNA and RNA from the patient sample as well as two controls, NA12878 and R403Q iPSC-CM, cells from a patient with HCM caused by a point mutation rather than a splice-site variant [Figure 1b].

**Figure 3:**
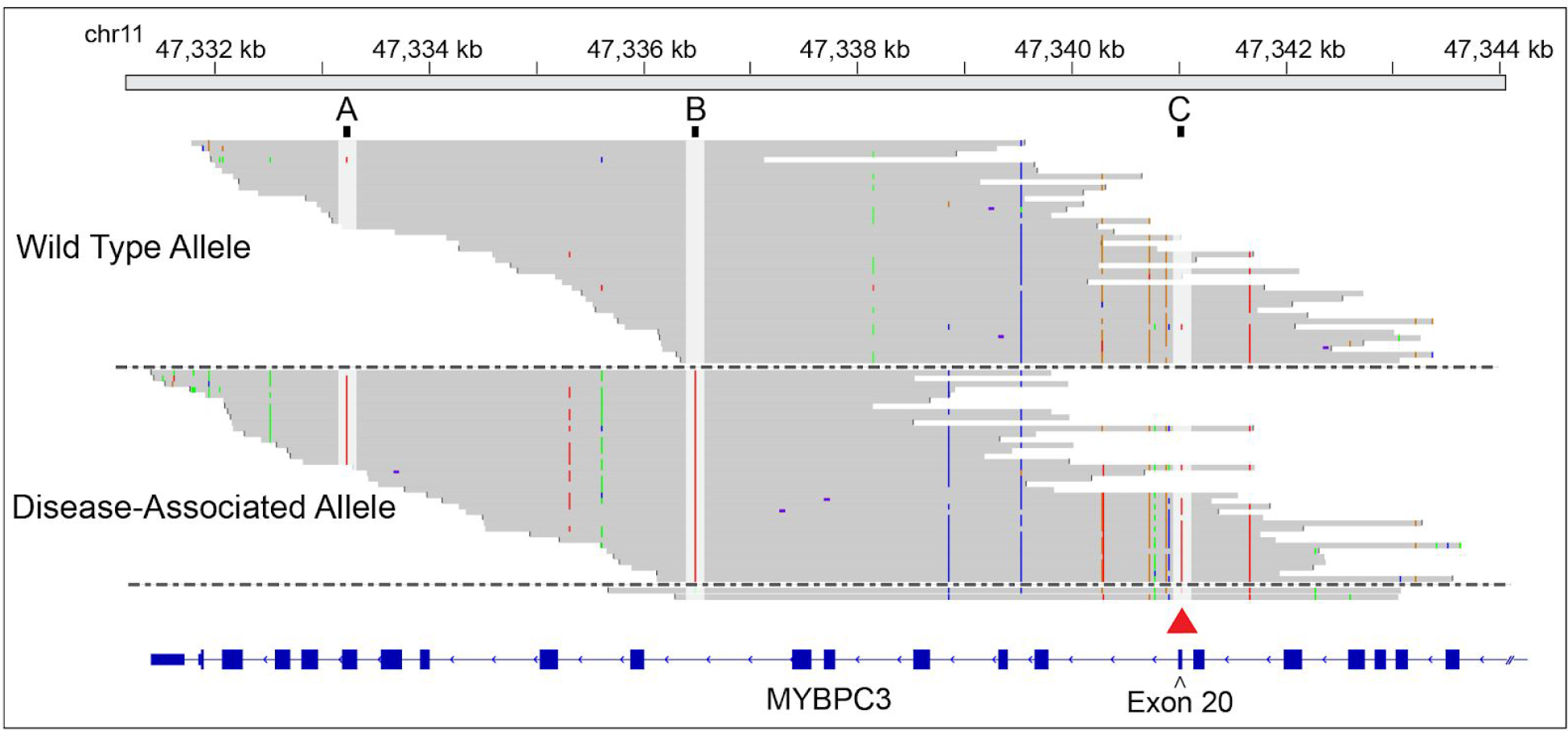
Targeted Long-Read Genome Sequencing For Phasing of Intronic and Exonic Variants. Long-read sequencing of target regions allowed for straightforward phasing of genomic regions. Here we show aligned reads from PacBio sequencing of targeted reads from the study patient, mapped to hg38 using minimap2. Reads were grouped based on the presence or absence of an alternate base at an intronic variant (rs11039189, chr11:47,336,484, here position B). Reads not covering this base were removed for clarity. This allowed us to easily sort reads into the wild type or alternative allele based on the presence or absence of the splice site variant in either of these alleles at position C (also marked with red triangle). We were also able to assign the alternate base at an exonic SNP (rs1052373, chr11:47,333,236, here position A) to the disease-associated allele. This allowed us to easily assign alternatively spliced isoforms in our transcriptome sequencing to either the wild type or disease-associated allele determined by the base called at rs1052372.

### PacBio Sequencing of Cardiac Gene Panel

Genomic sequencing coverage varied gene to gene, with TTN showing both highest coverage measured by read depth and percentage of the gene covered (1388 maximum read depth and >99.9% coverage of gene at least 1X) while *MYBPC3* showed the lowest maximum read depth (41 reads) and RYR2 showed the lowest total percentage of gene covered (59%) [NA12878, Supplemental Figure 4b]. Transcriptomic coverage also varied by genes, with TTN again showing the highest maximum read depth (11374X) but with multiple genes (*MYBPC3*, *MYH6*, *MYH7*, *TNNT2*, *TTN*) showing 100% coverage of exonic loci at ≥ 20X [Patient Sample, Supplemental Figure 5b].

### MinION Sequencing of Cardiac Gene Panel

Genomic sequencing coverage varied gene to gene, with TTN showing both highest coverage measured by read depth and percentage of the gene covered (891 maximum read depth and >99.9% coverage of gene at at least 1X) while *MYBPC3* showed the lowest maximum read depth (26 reads) and RYR2 showed the lowest percentage of gene covered (61%) [NA12878, Supplemental Figure 4a]. Transcriptomic coverage also varied by genes, with TTN again showing the highest maximum read depth (287X) but with multiple genes (*MYBPC3*, *MYH6*, *MYH7*, *MYL2*, *TNNT2*, *TTN*) showing 100% coverage of exonic loci at ≥ 1X [Patient Sample, Supplemental Figure 4b]. While depth is lower in our MinION sequencing of cDNA libraries, this is likely due to the comparatively small amount of cDNA spiked into the genomic pool.

### Phasing of Genomic Alleles

Targeted genomic reads from both technologies averaged around 7.5kb in length. This allowed us to easily phase a well-covered region of *MYBPC3* in our patient that contained both the splice-site variant and an exonic variant. [Figure 3] This information allowed us to later assign our alternatively spliced isoforms to either the wild-type or variant-containing allele based on this benign SNP.

### Unexpected Alternative Isoforms

Alignment of transcriptomic reads from both PacBio and Oxford Nanopore sequencing of our patient sample revealed alternatively spliced isoforms of *MYBPC3* [Figure 4a] in the region of the splice-site variant that included the expected and previously seen loss of exon 20, but also included additional unexpected isoforms. These included isoforms that retained the intron between exon 19 and exon 20, isoforms that used an alternate splice-donor site for exon 20, and some that included a cryptic exon between exons 20 and 21 [Figure 4b]. All of the new exon start and end sites contained proper splice-site sequences, with the exception of AS 1.5 and 1.6, where the extended exon 20 splice donor site contained a GC rather than a GT. The alternative splicing patterns seen in AS 1.2-1.10 resulted in the inclusion of premature stop codons. These would therefore not produce full length protein and are likely targeted for nonsense mediated decay. Two of these additional alternative splicing patterns (AS 1.2 and AS 1.8, Figure 4b) were seen in our original, primer-based IsoSeq data when we returned to examine that dataset (Supplemental Figure 1]. By aligning reads from each sample to our identified exon 19-20 alternative splicing patterns, we were able to assign them to either the wild-type or variant allele, and show that the alternatively spliced isoforms originated predominantly from the variant-containing allele [Figure 5b].

**Figure 4:**
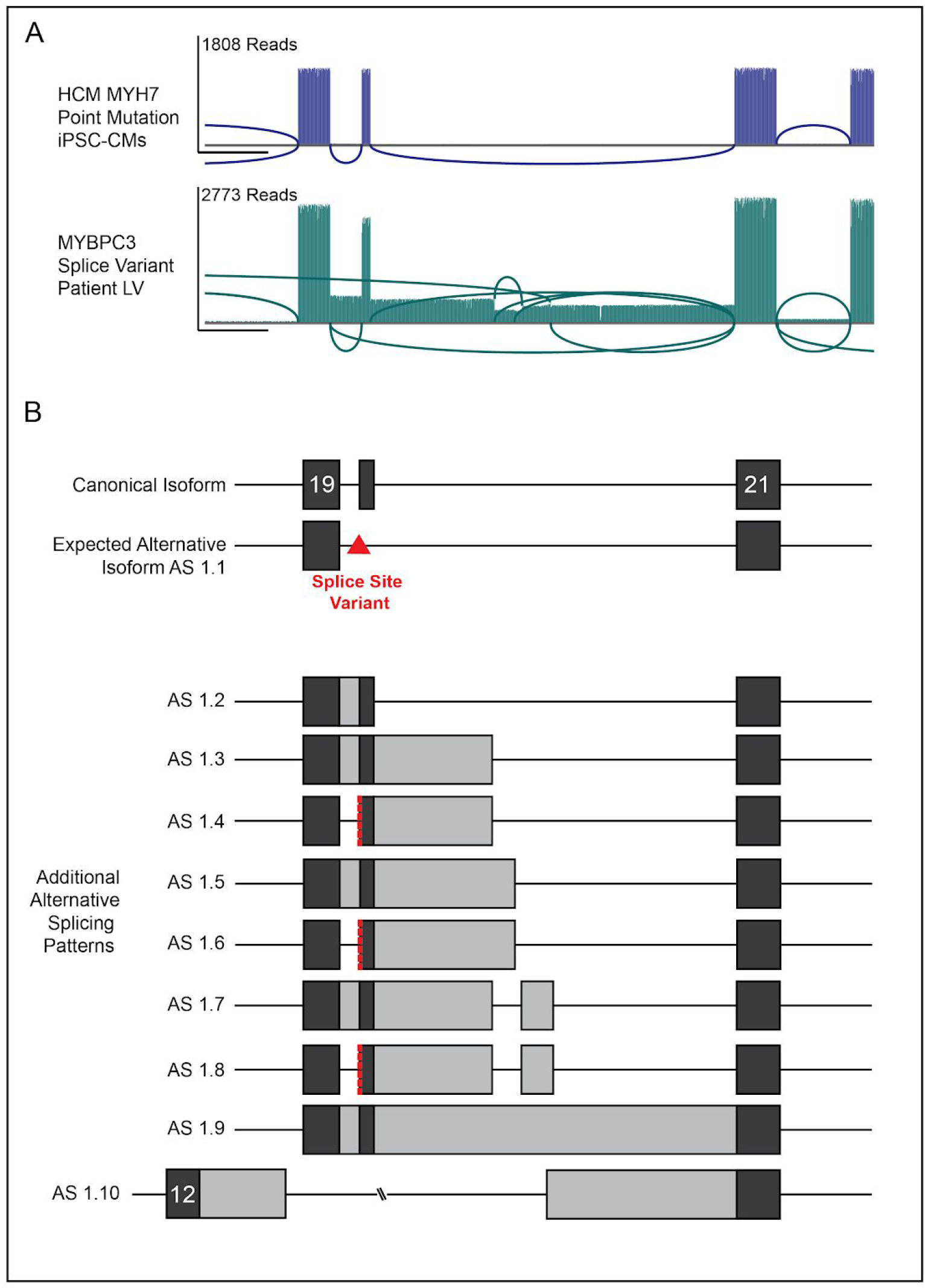
Alternative Splicing Patterns of *MYBPC3* in HCM Patient Sample. Sequencing of pulldown cDNA from the patient sample revealed not only expected wild-type and exon-20-missing (AS 1.1) isoforms but also additional alternative splicing events. Sashimi plots in panel A show a single splicing path through exons 19, 20, and 21 of *MYBPC3* in our HCM iPSC-CMs containing an *MYH7* point mutation while our *MYBPC3* splice-variant patient shows an abundance of alternative splicing junctions. The plots represented here are from PacBio sequencing of cDNA from the HCM iPSC-CMs and the first of two RNA extractions from the *MYBPC3* splice-variant patient. Panel B represents cartoon diagrams of the alternative splicing patterns observed in this region. Each splicing pattern was manually identified in IGV and used to build a reference file of alternatively spliced isoforms, the canonical isoform, and truncated, early-terminating isoforms found in both our patient and HCM control samples. For the purpose of investigating effects of the splice-site mutation on the exon 19-21 region, alternative splicing upstream and downstream of this region found in both the control HCM sample and the patient sample was ignored for this analysis. Splicing patterns AS 1.4, AS 1.6, and AS 1.8 use a slightly shifted splice acceptor site for exon 20, two bases downstream of the canonical site, denoted here by the red dashed line.

**Figure 5:**
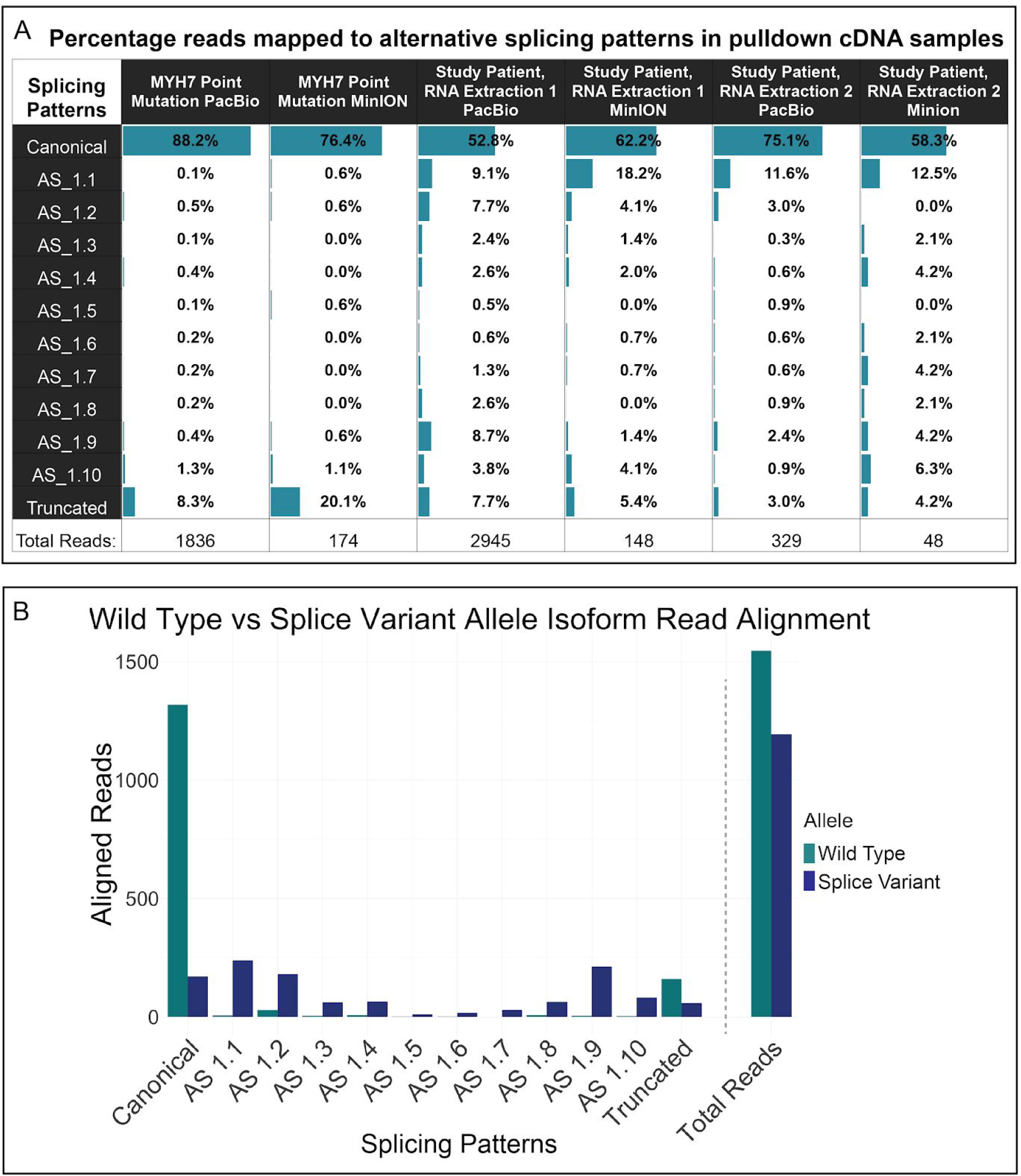
Quantification of alternatively spliced reads across samples and alleles. Demultiplexed sequencing data was remapped to the list of isoforms using minimap2 to analyze prevalence of each alternative splicing pattern in the sample. Panel A shows the prevalence of each splicing pattern across six cDNA pulldown sequencing samples: PacBio and MinION sequencing of our iPSC-CM sample as well as two unique RNA extractions from our *MYBPC3* splice variant patient sample. The alternatively spliced transcripts that appear at low percentages in our *MYH7* point mutation sample are predominantly results of mismapping when mapping such closely related reads, and stand as an example of background noise against which to compare the *MYBPC3* splice-variant alignments. Manual phasing in IGV using reads with a base called at rs1052373 (or rs10769255 for the truncated isoforms) revealed that the alternatively spliced transcripts were overwhelmingly generated from the splice-site variant containing alleles. Panel B shows representative phasing of reads mapped to each allele for PacBio sequencing of the first of two RNA extractions from our patient sample. Reads that contained neither the reference nor the known SNP base at rs1052373 or rs10769255 (likely due to inherent sequencing errors, and accounting for only 8.4% of total reads) were not included in this analysis. Here again, canonically spliced transcripts that arise from the mutant allele are in some part likely the result of mismapping against closely related isoforms and the intrinsic low level base calling errors found in long-read sequencing.

## DISCUSSION

Clinical genome sequencing is fast becoming a staple of modern medical care, allowing physicians to more precisely diagnose, treat, and manage care for patients with diverse diseases, including cardiovascular diseases. Panel based screening, exome sequencing, and whole genome sequencing all have the capacity to reveal disease-associated genetic variants. Yet in cases where those variants are novel or rare, and where definitive disease-causality cannot be determined, sequencing of RNA may be an important strategy for providing additional molecular evidence of disease-association. We here use targeted genomic and transcriptomic sequencing to link alternatively spliced isoforms of *MYBPC3* to a suspected disease-causing variant in a patient with HCM. Our sequencing revealed not only a predicted alternative isoform but also multiple additional novel isoforms resulting from the mutant allele. These isoforms provide strong additional evidence of disease-causality for this variant, which can allow for more directed treatment and management of disease in not only this patient but also potentially affected family members.

We used two strategies for targeting disease-associated genes and transcripts in this study. Our first targeting method used gene-specific primers to amplify *MYBPC3* cDNA from ten left-ventricular heart samples, including six controls and four HCM patients. Isoform analysis performed using PacBio’s IsoSeq platform revealed highly-expressed alternatively-spliced isoforms in a patient with a predicted splice-site altering variant in *MYBPC3*. As this variant occurred in the splice-acceptor site preceding exon 20, we anticipated that transcripts generated by the mutant allele may be missing this exon. Indeed, our primer-based targeting revealed that while the most highly expressed isoform of *MYBPC3* in this sample was the canonical isoform, the next most highly expressed isoform was missing exon 20 [Figure 2].

We proceeded to perform additional targeted sequencing using a panel of commercially available oligonucleotide probes designed to target 10 cardiac genes of interest. We used these probes to selectively pulldown fragmented genomic DNA as well as cDNA from our *MYBPC3* splice-site variant patient, iPSC-cardiomyocytes derived from an HCM patient with a missense mutation in *MYH7*, and NA12878. We demonstrated that these pulldown libraries made with commercially available probes could be sequenced not only with Pacific Biosciences SMRT sequencing but also with Oxford Nanopore Technologies MinION sequencing.

Our genomic pulldown libraries did not show uniform high depth coverage across all of our genomic loci. This is not unexpected, as many of these genes contain large introns which were not targeted by our exonic probe sequences. Other sources of uneven read depth are likely due to areas of close sequence similarity in gene family members (such as additional members of the myosin gene family, which have high sequence similarity to *MYH6* and *MYH7*) where probes may indiscriminately pull down off-target family members, reducing the number of probes available for on-target loci pulldown. Despite these potential issues with using exonic probes for genomic pulldown, we were able to achieve high enough coverage of *MYBPC3* on both PacBio and MinION platforms to confidently phase the intronic splice-site variant with an exonic SNP. Future targeted sequencing focused on fully and evenly covering all of these genes could include genomic probes spiked into regions of low coverage to fill in gaps.

Long-read transcriptomic sequencing allows us to look at full-length spliced transcripts, not just the prevalence of alternatively spliced junctions. This allowed us to not only phase the intronic *MYBPC3* variant with the expected alternatively spliced isoform, but also to identify additional alternative spilicng patterns generated by the mutant allele. These additional splicing patterns featured retained introns, extended exons, and the appearance in some isoforms of an additional cryptic exon [Figure 4b]. The alternative splicing patterns that retain what appears to be normal splicing at the splice acceptor site at exon 20 (AS 1.4, 1.6, and 1.8) actually make use of an alternative splicing site two bases downstream of the canonical exon 20 splice acceptor site. The mutation, which results in the destruction of the canonical site, therefore results in splicing at this slightly shifted site, resulting in an overall frameshift.

While the expected alternatively spliced isoform (AS 1.1, which resulted in the exclusion of exon 20) remained in frame, the additional splicing patterns all resulted in premature stop codons, likely targeting them for nonsense mediated decay. AS 1.1 retained the open reading frame and therefore may be translated into protein. As the deleted exon fell in a poorly defined linker region, it is hard to know what effect this may have had on protein function^27^. The shortened protein may have difficulty associating with its appropriate binding partners in the cardiac sarcomere, or may interfere with normal contractile motion of the sarcomere. The additional isoforms containing premature stop codons may be targeted for nonsense mediated decay, potentially lowering overall levels of *MYBPC3* RNA and protein expression. Research has shown that haploinsufficiency of *MYBPC3* during differentiation can lead to HCM-like phenotypes in a cell model of disease^15^. Recent work has also suggested that reduced levels of *MYBPC3* result in increased sarcomere contractility^28^. Both shortened isoforms missing exon 20 and a reduction in the overall amount of expressed *MYBPC3* may therefore have contributed to disease in this patient. Further studies of the effect of the loss of exon 20 on *MYBPC3* protein function may shed additional light on not only how this isoform may cause disease but also the role that this small linker region plays in *MYPBC3* function.

Clinical genome sequencing can allow for better diagnosis, treatment, and management of disease. Yet genome sequencing alone may not be enough to provide a causal link between variant and disease in the case of rare or de novo variants. We suggest that targeted, long-read RNA sequencing in conjunction with genome sequencing may provide additional molecular evidence of disease for these variants, as well as providing new information about the consequence of these variants on downstream RNA and protein expression. Targeted sequencing of both the genome and the transcriptome in our patient sample revealed transcript level consequences of a rare, uncharacterized variant in a patient with HCM. Using long-read sequencing rather than traditional short read sequencing allowed us to easily characterize alternatively spliced isoforms and to assign them to the variant-associated allele, providing additional evidence of disease-causality. This strategy may prove to be effective in providing additional molecular evidence of disease-causality for rare or de novo variants not only in patients with HCM but in many diseases.

