## Supplemental Material for "Targeted Long-Read RNA Sequencing Demonstrates Transcriptional Diversity Driven by Splice-Site Variation in *MYBPC3*"

| Gene | Primer Set | FWD | REV |
|---|---|---|---|
| MYBPC3 | Set 1 | /5Phos/tcagacgatgcgtcatAGTCCCTCTTTGGGTGACCT | /5Phos/atgacgcatcgtctgaGCCCAATAAACATTGGGAAG |
| MYBPC3 | Set 2 | /5Phos/ctatacatgactctgcAGTCCCTCTTTGGGTGACCT | /5Phos/gcagagtcatgtatagGCCCAATAAACATTGGGAAG |
| MYBPC3 | Set 3 | /5Phos/tactagagtagcactcAGTCCCTCTTTGGGTGACCT | /5Phos/gagtgctactctagtaGCCCAATAAACATTGGGAAG |
| MYBPC3 | Set 4 | /5Phos/tgtgtatcagtacatgAGTCCCTCTTTGGGTGACCT | /5Phos/catgtactgatacacaGCCCAATAAACATTGGGAAG |
| MYBPC3 | Set 5 | /5Phos/gatctctactatatgcAGTCCCTCTTTGGGTGACCT | /5Phos/gcatatagtagagatcGCCCAATAAACATTGGGAAG |

**Supplemental Table 1: Gene Specific, Barcoded Primers for preliminary Iso-Seq of *MYBPC3***

Primers were designed to the first and last exons of *MYBPC3*. These sequences (capital letters) were maintained between primers, but individual barcodes (lowercase letters) were designed for each set, such that pooled samples could later be identified.

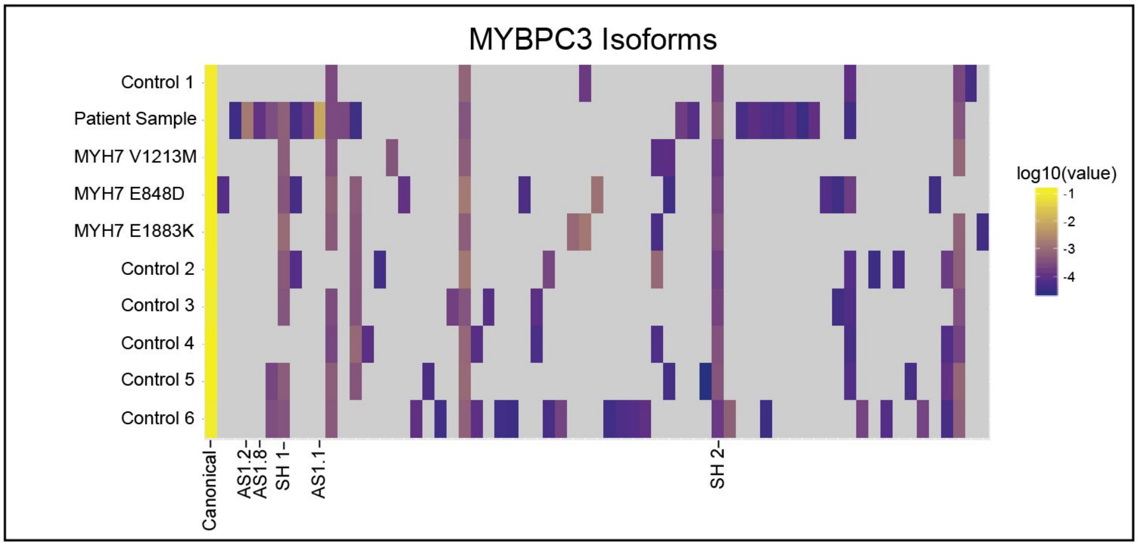


**Supplemental Figure 1: Chaining of all *MYBPC3* Isoforms from preliminary Iso-Seq experiments reveals one dominant isoform per sample** Chaining of *MYH7* and *MYBPC3* isoforms across samples revealed one dominant isoform (first column, yellow) across all samples. However one sample (Patient Sample) showed an increased number of highly expressed *MYBPC3* alternative isoforms as compared to the other ten samples in panel A. This sample also showed one highly expressed alternate isoform (AS 1.1). The six labelled isoforms correspond to the isoforms depicted in Figure 2 panel A as well as AS 1.8, also described in Figure 4 panel B.

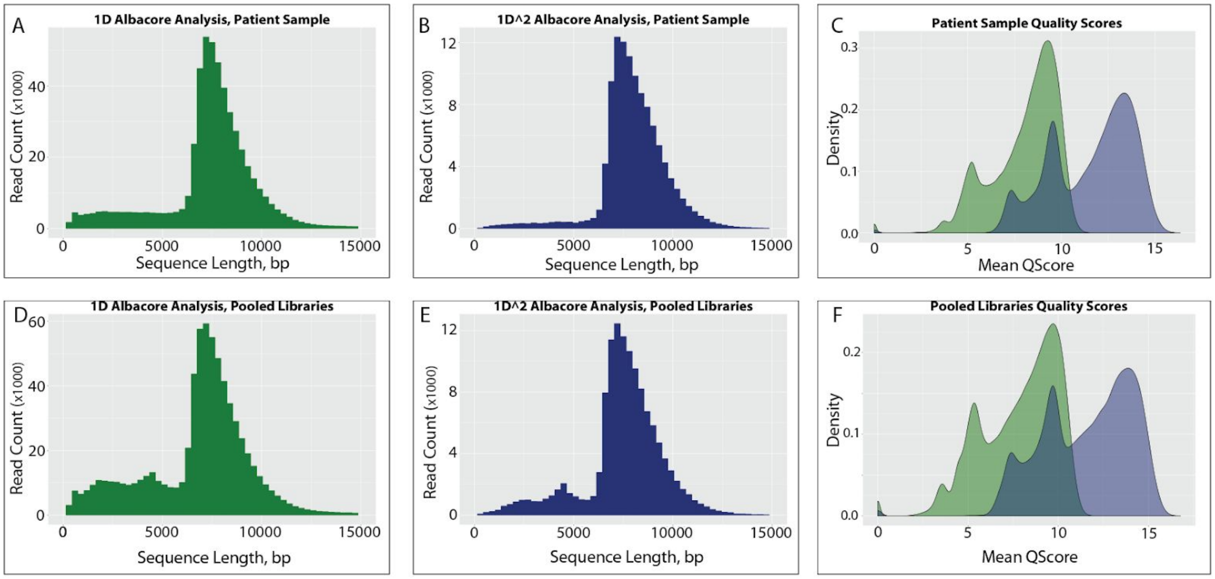


G

| Sample | Seq Type | Mean Length (bp) | Median Length (bp) | Read Count | Total Sum (bp) | Pass Quality Cutoff (bp) |
|---|---|---|---|---|---|---|
| Patient Sample | 1D | 7425 | 7598 | 527892 | 3919678682 | 2970632384 |
| Patient Sample | 1D^2 | 7998 | 7927 | 113126 | 904873660 | 562613389 |
| Pooled Libraries | 1D | 6836 | 7235 | 702719 | 4804089848 | 3350280515 |
| Pooled Libraries | 1D^2 | 7465 | 7593 | 126296 | 942840393 | 563379905 |

**Supplemental Figure 2: Sequence statistics from MinION R9.5 Flowcells.** Two MinION R9.5 flowcells were used for 1D^2 sequencing: one for genomic pulldown from the patient sample (a-c) and one for pooled genomic and cDNA pulldowns from all three samples (d-f). Output reads were basecalled using Albacore v2.2.7, first for 1D calls of each read (a,d) and then for paired 1D^2 reads (b,e). Sequence length distributions for the genomic pulldown alone fell between our library size selection of 6-13kb for both 1D and 1D^2 reads (a,b), while the pooled genomic and cDNA library showed an increase in shorter fragments, representing our 3-9kb size selected cDNA population which accounted for about 10% of the pool (d,e). As anticipated, read qualities of both libraries increased between 1D analysis (green) and 1D^2 analysis (blue, c,f). The table in G summarizes overall statistics for each, including patient sample, sequencing analysis type (Seq Type), mean and median read lengths, read count, total sequencing sum, and the sum of sequencing reads that passed Albacore's intrinsic quality cutoff scores.

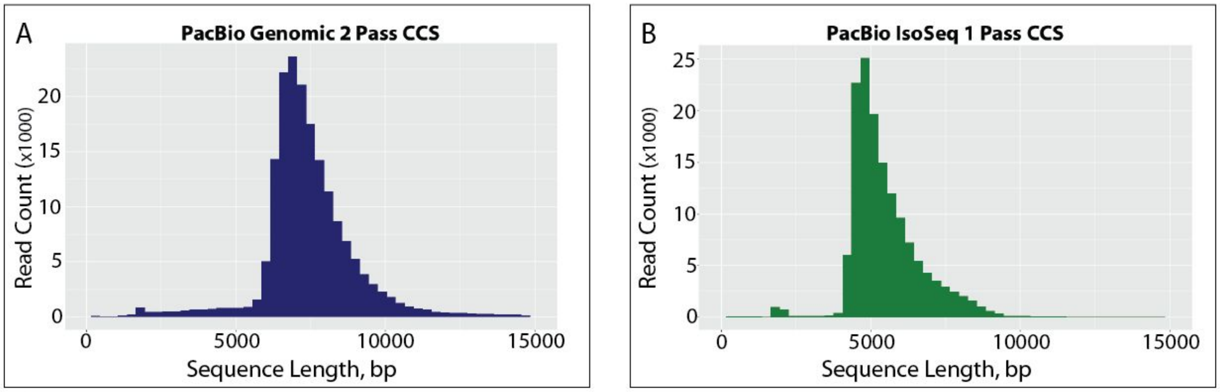


C

| Sample | CCS # | Mean Length (bp) | Median Length (bp) | Read Count | Total Sum (bp) | Total SMRTCell Sum (bp) |
|---|---|---|---|---|---|---|
| Pooled Genomic | 2 | 7465 | 7241 | 182172 | 1360081630 | 13419515077 |
| Pooled IsoSeq | 1 | 5507 | 5189 | 146522 | 806947849 | 8836246298 |

**Supplemental Figure 3: Sequencing Statistics from Sequel SMRT cells.** Two Sequel SMRT cells were used for sequencing, one for pooled genomic pulldown of all three samples and one for pooled cDNA pulldowns from all three samples. Plotted are read length distributions for the genomic pulldown pool after calling CCS (circular consensus sequence) for reads that had at least two full reads of insert (a) and the cDNA pulldown pool after calling CCS for reads that had at least 1 full read of insert (b). Sequence length distributions for the genomic pulldown fell between our library size selection of 6-13kb (a), while the pooled cDNA library showed an increase in shorter fragments, representing our 3-9kb size selected cDNA population (b). Unlike the MinION reads, Sequel data does not provide per-base or per-read quality scores. The table in C summarizes overall statistics for each, including the type of sample sequence, number of reads of insert required for CCS, mean and median read length, total read count, the total sum of CCS read sequence, and the total sum of sequence export from each SMRT cell before CCS calling.

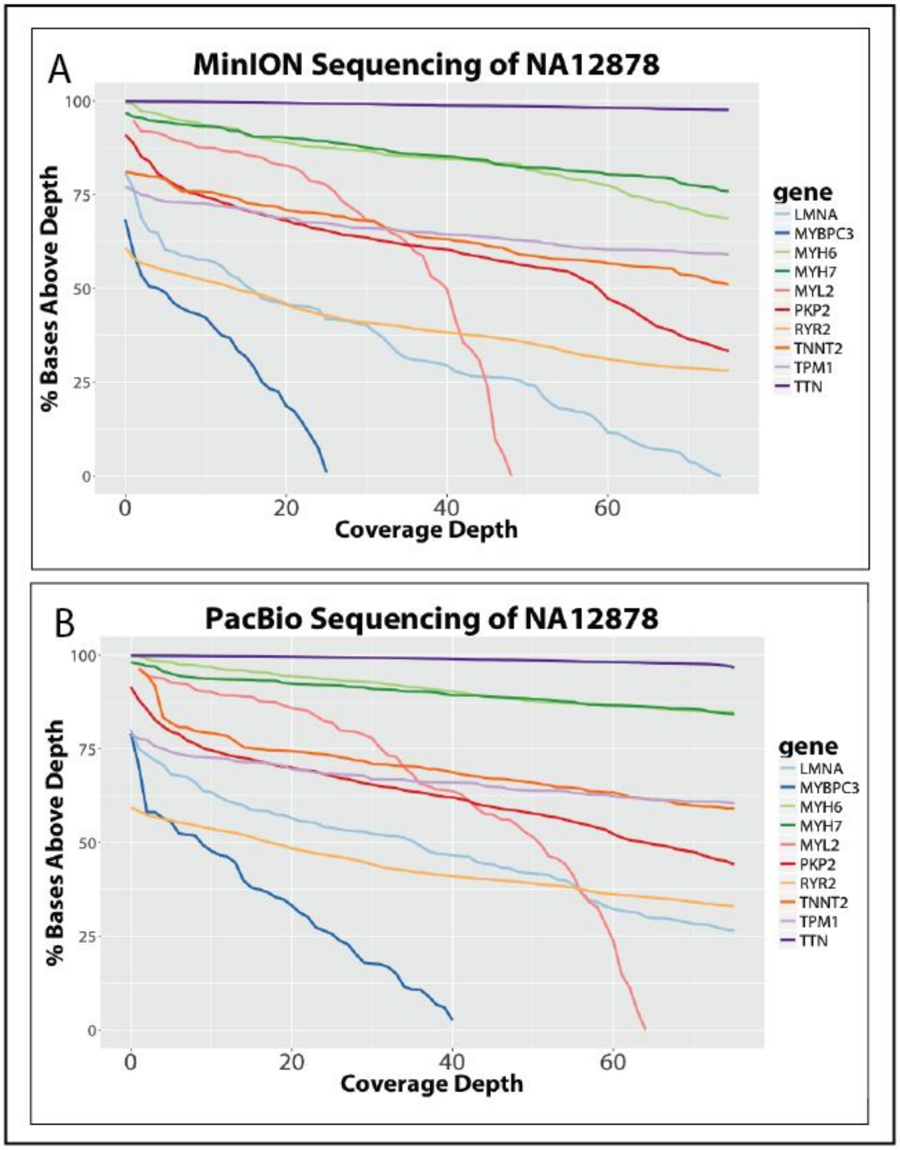


**Supplemental Figure 4: Genomic Sequencing Coverage Across 10 Pulldown Pool Genes in NA12878.** Coverage of either MinION (a) or PacBio (b) sequencing of 10 cardiac genes of interest after pulldown and pooled sequencing in NA12878. MinION sequencing shown is 1D^2 sequencing while PacBio sequencing is genomic sequencing with 2 reads of insert required for CCS. Both samples were mapped with minimap2 to hg38. Shown here are coverages up to 75X, though some samples extend to higher coverages, including areas of TTN reaching a maximum depth of 891 after MinION sequencing and 1388 after PacBio sequencing. Uneven coverage across each gene is likely due to the nature of the pulldown probes, which were designed to target exons rather than the complete genomic regions. Gene loci specified from NCBI RefSeq coordinates.

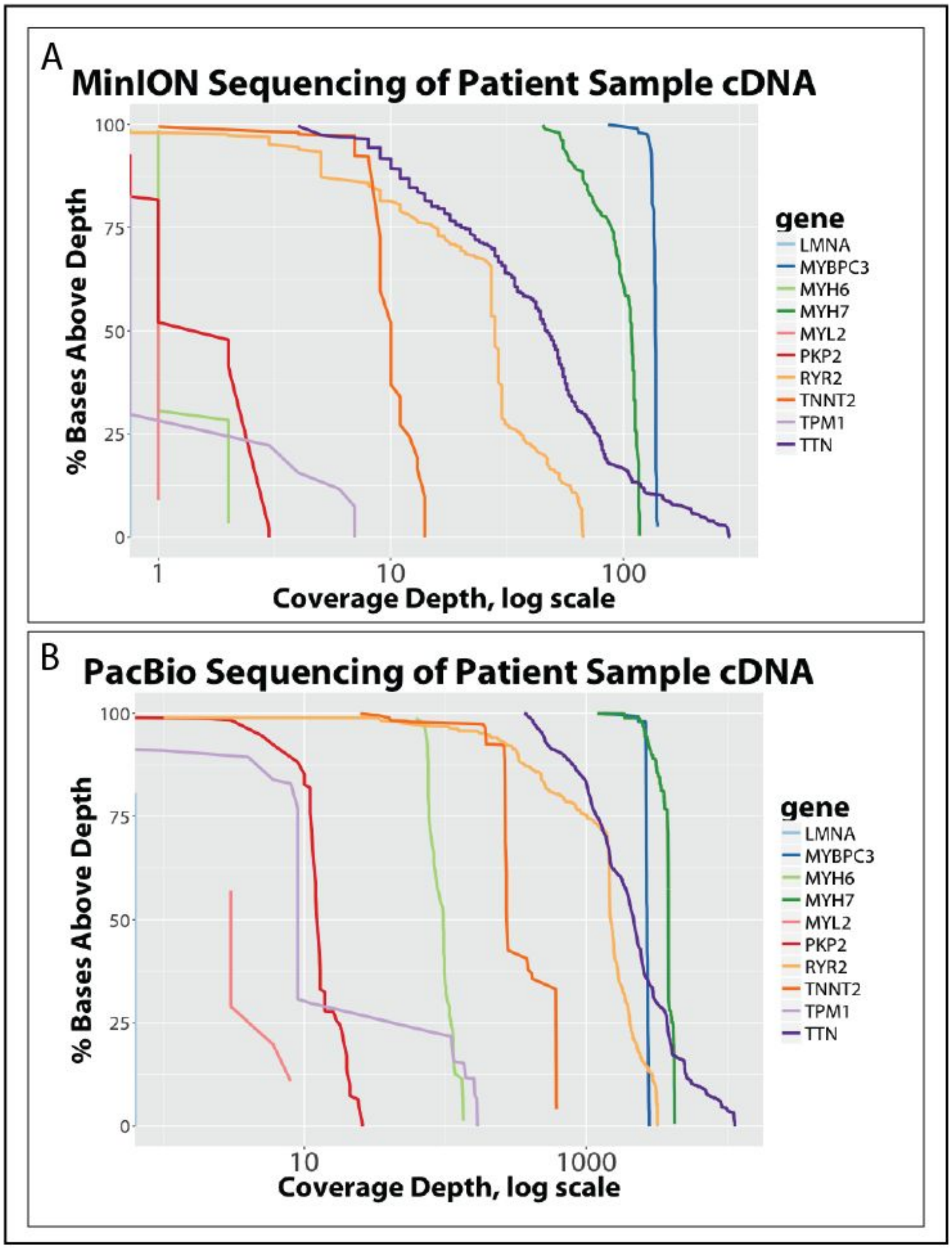


**Supplemental Figure 5: Transcriptomic Sequencing Coverage Across 10 Pulldown Pool Genes in Patient Sample.** Coverage of either MinION (a) or PacBio (b) cDNA sequencing of 10 cardiac genes of interest after pulldown and pooled sequencing in our MYBPC3 splice-site variant patient. These coverage plots represent a single cDNA library from the first of two RNA extractions from this patient. MinION sequencing shown is 1D^2 sequencing while PacBio sequencing is cDNA sequencing with 1 read of insert required for CCS. Both samples were mapped with minimap2 to hg38. Coverage depth varies across genes, with highly expressed genes like *MYH7* and MYBPC3 covered at greater than 45X (*MYH7*) and 86X (MYBPC3) in MinION sequencing and 1199X (*MYH7*) and 1304X (MYBPC3) across all exons in PacBio sequencing. Exonic loci specified from NCBI RefSeq coordinates.

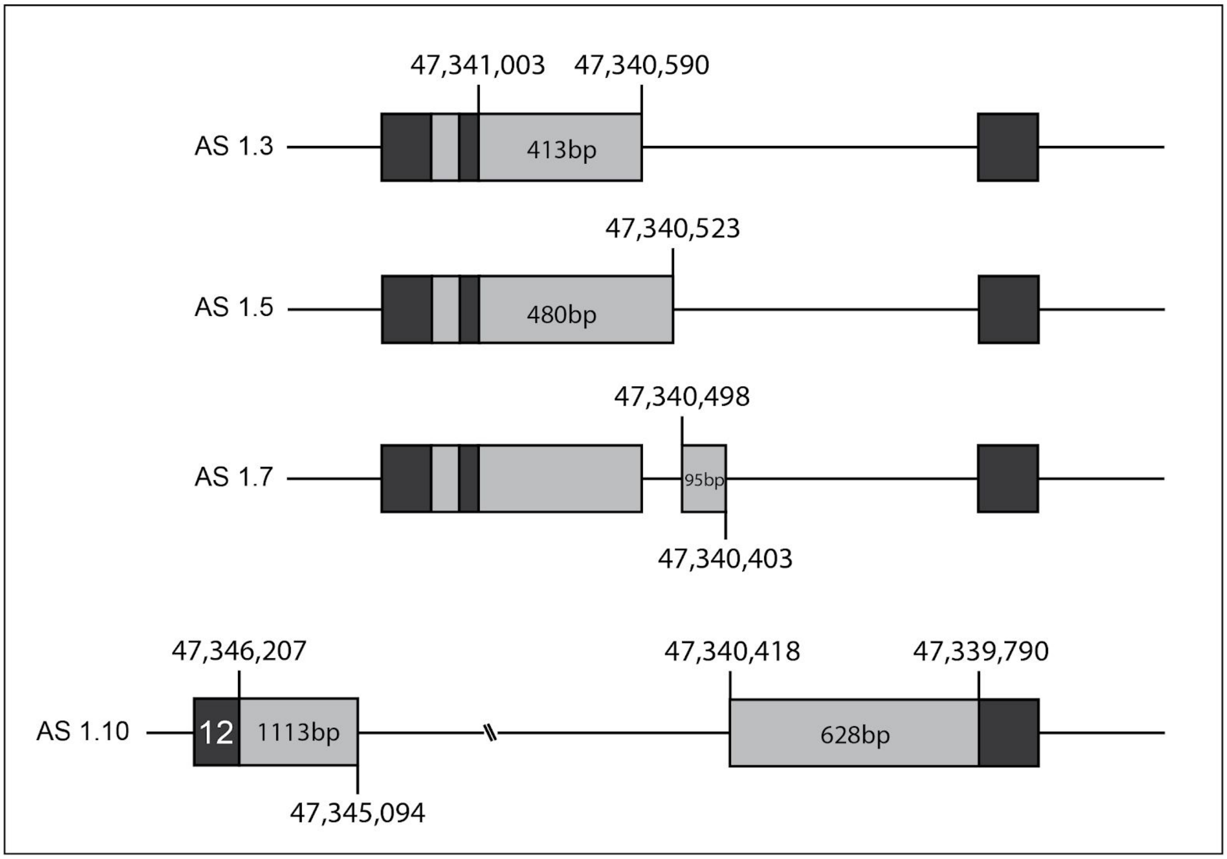


**Supplemental Figure 6: Start and end points of non-canonical splice sites in *MYBPC3*** Displayed are the start and end points of the alternative splicing patterns for exon-extension or pseudo-exon events described in Figure 4, as demonstrated on the first listed isoform in which each appears. Also displayed are the lengths in bp of these exon extensions. The exon 12 extension event in AS 1.10 is not displayed to scale. There was slight variation in the AS 1.10 junction which was potentially due to mismapping. Alignments were performed against the predominant junction shown here. Locations displayed are hg38, chromosome 11.
